# Sustained attention as measured by RT variability is a strong modulator for the P600, but not the N400

**DOI:** 10.1101/2021.11.18.469143

**Authors:** Friederike Contier, Mathias Weymar, Isabell Wartenburger, Milena Rabovsky

**Affiliations:** Cognitive Sciences, University of Potsdam, Germany

**Author notes:** Corresponding author: Friederike Contier, Cognitive Neuroscience Lab, Department of Psychology, University of Potsdam, Karl-Liebknecht-Str. 24-25, 14476 Potsdam, Germany.

**Keywords:** P600, N400, sustained attention, automaticity vs control, reaction time variability

## Abstract

The functional significance of the two prominent language-related ERP components N400 and P600 is still under debate. It has recently been suggested that one important dimension along which the two vary is in terms of automaticity versus attentional control, with N400 amplitudes reflecting more automatic and P600 amplitudes reflecting more controlled aspects of sentence comprehension. The availability of executive resources necessary for controlled processes depends on sustained attention, which fluctuates over time. Here, we thus tested whether P600 and N400 amplitudes depend on the level of sustained attention. We re-analyzed EEG and behavioral data from a sentence processing task by Sassenhagen & Bornkessel-Schlesewsky (2015, *Cortex*), which included sentences with morphosyntactic and semantic violations. Participants read sentences phrase by phrase and indicated whether a sentence contained any type of anomaly as soon as they had the relevant information. To quantify the varying degree of sustained attention, we extracted a moving reaction time coefficient of variation over the entire course of the task. We found that the P600 amplitude was significantly larger during periods of low reaction time variability (high sustained attention) than in periods of high reaction time variability (low sustained attention). In contrast, the amplitude of the N400 was not affected by reaction time variability. These results thus suggest that the P600 component is sensitive to sustained attention while the N400 component is not, which provides independent evidence for accounts suggesting that P600 amplitudes reflect more controlled and N400 amplitudes more automatic aspects of sentence comprehension.

---

Event-related potential (ERP) components observed during sentence processing play an important role for neurocognitive models of language comprehension (e.g., Bornkessel-Schlesewsky & Schlesewsky, 2013; Brouwer et al., 2017; Friederici, 2011; Kuperberg, 2007, 2021; Rabovsky et al., 2018). The N400 is mostly sensitive to the semantic fit of a word within a given context, with more negative amplitudes the less predictable the word (Kutas & Hillyard, 1980). The later P600 is triggered by a wide range of syntactic violations, semantic incongruencies, syntactic ambiguities and even pragmatic factors and spelling errors (e.g., Hagoort et al., 1993; Kim & Osterhout, 2005; Osterhout & Holcomb, 1992; Regel et al., 2014; Van de Meerendonk et al., 2011). However, even after four decades of empirical studies, their functional significance is still actively debated (see, e.g., discussions in Kutas & Federmeier, 2011; Leckey & Federmeier, 2019).

Recently, it has been suggested that the two components differ along the automaticity vs control dimension (e.g., Kolk & Chwilla, 2007; Kuperberg, 2007; Rabovsky & McClelland, 2020; van Gaal et al., 2014). This distinction and related dichotomies, such as between conscious vs unconscious processes, are ubiquitous across domains in cognitive (neuro)science (Posner & Snyder, 1975; Schneider & Shiffrin, 1977). Language comprehension often seems like a fast and effortless process and indeed, N400 amplitude might reflect a more automatic brain response, for instance signaling the change in a probabilistic representation of sentence meaning corresponding to an implicit semantic prediction error (Rabovsky & McClelland, 2020). Sometimes however, this automatic process might not result in a coherent interpretation and an additional, more controlled process might be needed for successful comprehension. This process could be reflected in the P600 as a controlled revision process when automatic update fails, resulting in an initial state of uncertainty or conflict (Rabovsky & McClelland, 2020). Relatedly, the positivity might act as an error monitoring signal, where the language system interacts with the domain-general executive system (Kolk et al., 2003; Kolk & Chwilla, 2007; Van De Meerendonk et al., 2010).

There are several lines of evidence supporting the automaticity vs control distinction regarding the N400 and P600. For example, the P600 – but not the N400 – is modulated by task relevance, with a reduced or absent P600 when the instructions do not require the participant to process the anomaly (Gunter & Friederici, 1999; Hahne & Friederici, 1999; Molinaro et al., 2011; Schacht et al., 2014; Vissers et al., 2007) or participants do not notice the anomaly (Batterink & Neville, 2013; Osterhout & Mobley, 1995; Xu et al., 2019). The two components are also differentially affected by error probability, with increasing number of violations within an experimental stimulus set diminishing the P600, but not affecting the N400 (Coulson et al., 1998; Hahne & Friederici, 1999; Yano et al., 2021). Relatedly, the P600, but not N400, is absent during the attentional blink and other manipulations testing conscious vs unconscious processing of linguistics stimuli (Batterink & Neville, 2013; Kiefer, 2002; Luck et al., 1996; Rohaut & Naccache, 2017; Service et al., 2007; van Gaal et al., 2014). The P600 further correlates with indices of executive control (Brothers et al., 2021), exhibits sequential adaptation effects (Xu et al., 2021), and relates to eye movement regressions during natural reading (Dimigen et al., 2007; Metzner et al., 2017). Lastly, the P600 has also been linked to the more domain-general P3 component (Coulson et al., 1998; Sassenhagen et al., 2014; Sassenhagen & Bornkessel-Schlesewsky, 2015; Sassenhagen & Fiebach, 2019), which has been related to, for instance, stimulus saliency, surprise, and context updating (Donchin, 1981; Nieuwenhuis et al., 2005; Polich, 2007). Despite this previous evidence, the proposed distinction between the P600 and N400 is still debated. Some argue that the process underlying the P600 might also reflect a default process, for instance sentence meaning integration (Brouwer et al., 2017). Conversely, the N400 might also involve controlled aspects of sentence comprehension (Batterink et al., 2010; Lau et al., 2008) or it might not be possible to clearly categorize the component within this dichotomy (Kutas & Federmeier, 2011).

Generally, the availability of executive resources necessary for controlled processes - but less required for automatic processes - depends on the level of sustained attention. Sustained attention, the focus on a task over a period of time which is also known as “vigilance”, is enabled by arousal (possibly via a noradrenergic route) and in turn, modulates the amount of cognitive resources and resulting performance (Esterman & Rothlein, 2019; Oken et al., 2006). When arousal is in an optimal, medium range during task engagement, sustained attention is high, so more cognitive resources are available, which leads to high performance (e.g., fast reaction times and low error rates). During very low or very high arousal, sustained attention to the current task is reduced (e.g., during mind wandering), resources for the task at hand are limited, and performance suffers. Indeed, the actual level of sustained attention correlates with, for instance, response inhibition (Bellgrove et al., 2004; Connolly et al., 2005; Esterman et al., 2013), set shifting (MacDonald et al., 2009), and selective attention (Weissman et al., 2006). Under current view, sustained attention is by no means an all-or-nothing phenomenon but fluctuates over time (e.g., over the course of a task; Esterman et al., 2013; Fortenbaugh et al., 2018; Johnson et al., 2015; Van Den Brink et al., 2016).

That attention enables executive resources has important implications for the processes underlying the P600 and N400. If the P600 indeed reflects a controlled process, it should be reduced when sustained attention is low, and thus, less of the necessary executive resources are available. Conversely, if the N400 reflects an automatic process, its amplitude should not depend on sustained attention. To date, direct evidence on the role of attention on the P600 and N400 besides the manipulation of task relevance (see above) is still scarce.

To further investigate whether the P600 and N400 might be dissociated on the automaticity versus control dimension, we thus tested their susceptibility to sustained attention in a sentence processing paradigm using reaction time variability as an index of sustained attention. We re-analyzed EEG and behavioral data from a visual sentence processing task by Sassenhagen & Bornkessel-Schlesewsky (2015, henceforth SBS). SBS investigated the relationship between reaction time measures and language-related ERPs and found that the P600 latency is aligned with the RT rather than stimulus onset. Sentences in their task were either correct, contained a morphosyntactic, or semantic violation. Instead of a typical grammaticality judgment prompt *following* each complete sentence, participants were instructed to respond whether a sentence contained any type of anomaly as soon as they had the relevant information. To continuously measure fluctuations in sustained attention, we focused on reaction time variability within participants. The underlying idea is that in periods when sustained attention is high, there is less variability in reaction times than when sustained attention is low. We quantified sustained attention with a moving reaction time coefficient of variation over the entire course of the task for each participant (see e.g., Esterman et al., 2013; Van Den Brink et al., 2016). We expect P600 amplitudes, but not N400 amplitudes, to be negatively predicted by RT variance, leading to larger amplitudes in periods of low RT variance (high sustained attention) than high RT variance (low sustained attention).

## Method

### Participants

Participants were right-handed, native speakers of German, with normal or corrected-to-normal vision, no history of neurological disorders, between the ages of 18 and 40, of either sex. For our analyses, we analyzed the data from the 20 participants that were also analyzed and made publicly available by SBS. This sample size is within the range where previous studies have found neural and pupillometric correlates of RT variability (Connolly et al., 2005; Esterman et al., 2013; Johnson et al., 2015; Van Den Brink et al., 2016). We report all data exclusions, all manipulations, and all measures in the study.

### Data Acquisition

Participants’ electroencephalogram (EEG) was recorded using a BrainProducts actiCHamp system (Brain Products GmbH, Gilching, Germany) at 1000 Hz, using 64 electrodes spaced according to the international 10-20 system. Impedances were reduced below 35 kOhm if possible. A forehead ground and a reference-free recording were used. Each subject took part in both a face detection experiment designed to elicit a P3 (not reported here), and a sentence processing experiment designed to elicit a P600 and N400. For more details on the data collection procedure, see SBS (p. A6).

### Sentence Processing Task

Participants read German sentences on screen phrase by phrase. Half of the sentences were control sentences which introduced a category word (hypernym) and three category members (hyponyms), for instance, “Zur Kategorie | Obst | gehören | der Apfel, | die Birne | und | die Mango” (lit: *To the category* | *fruit* | *belong* | *the apple*,| *the pear*,| *and* | *the mango*). In deviant sentences with a morphosyntactic violation, one of the hyponyms was preceded by a mismatching determiner regarding its grammatical gender (e.g., “das Birne”, *the*_neut_ *pear*_fem_). In semantically deviant sentences, one of the hyponyms was exchanged for a hyponym from another experimental sentence, and hence, another semantic category (e.g., “der Vogel”, *the bird*, instead of *the pear* in the example above). Single violations appeared equally often in position one, two, or three. A portion of the violation sentences contained a double violation, i.e., a morphosyntactic violation on one hyponym and a semantic violation on another (e.g., “Zur Kategorie | Obst | gehören | der Vogel, | das Birne | und | die Mango”, lit: *To the category* | *fruit* | *belong* | *the bird*_*(semantic violation)*_, | *the*_neut_ *pear*_fem *(morphosyntactic violation)*_, | *and* | *the mango*).

Stimulus presentation for each participant was randomized and comprised 150 correct control sentences, 75 sentences with a morphosyntactic as the only or first violation, and 75 sentences with a semantic violation as the only or first violation. Sentences appeared phrase by phrase, for 350 ms per phrase with a 350 ms blank screen between phrases (indicated by vertical lines in the example above). Importantly, participants were instructed to respond as quickly as possible whether a sentence was anomalous or not as soon as they had the relevant information. Specifically, they were instructed to “[…] press a pre-assigned button once they know the sentence to be either correct, or (structurally or semantically) incorrect.” (SBS, p. A7). Participants were encouraged to respond as quickly and accurately as possible by a) a specific feedback tone following correct and incorrect (or timeout) responses and b) displaying their average RT and accuracy in between trials. The negative feedback tone was also played following time-outs (responses > 2000 ms after last word). If accuracy dropped below 80% or mean RT rose above 1000 ms, they were additionally urged to respond more rapidly and accurately by the experimenter. For additional details on stimulus construction and the experimental procedure, see SBS (p. A6f.).

### Reaction Time Coefficient of Variation

To get a measure of sustained attention over the course of the task, we calculated how variable the RTs around each trial were, a method established as reaction time coefficient of variation (RT CV; Esterman et al., 2013; Van Den Brink et al., 2016). This is typically done by calculating the standard deviation over a number of trials which is a reasonable time period of an attentional state and dividing the result by the mean (normalization).

For each participant, we thus extracted RTs from every trial, that is, including both control and violation trials and irrespective of accuracy of the response. For violation trials, RT was time-locked to the onset of the respective violation phrase (in either position 1, 2 or 3). If the sentence contained a double violation, RT was time-locked to the onset of the first one. For control trials, RT was time-locked to the onset of the last phrase of the sentence. Unsurprisingly, RT data were not normally distributed (mean skewness across participants: 3.11), which necessitated log-transformation of the RTs. Conservatively, for trials in which participants responded before the onset of the respective phrase (mean = 9.1 trials, range = 2-19), the resulting negative values were replaced with NA before log-transformation, ignored for the subsequent calculation of the RT CV, and interpolated during the smoothing process in the end.

We then calculated the RT CV using an established CV formula specifically adjusted to log-normally distributed data (e.g., Julious & Debarnot, 2000). Specifically, for each trial, we calculated the RT CV value as 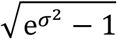, where *σ* equals the standard deviation of the log-transformed RTs in a given time period surrounding that trial. Our assumption was an attentional state of around 60 s and thus our planned window size over which each RT CV was calculated was 10 trials. However, to make sure our results are not conditional on this parameter choice, we additionally explored effects based on RT CVs over both a shorter (5 trials / ~30 s) and longer (15 trials / ~ 90 s) window. We then smoothed over the resulting RT CV line (see Fig. 1) using a Gaussian kernel of two trials.

**Figure 1.**
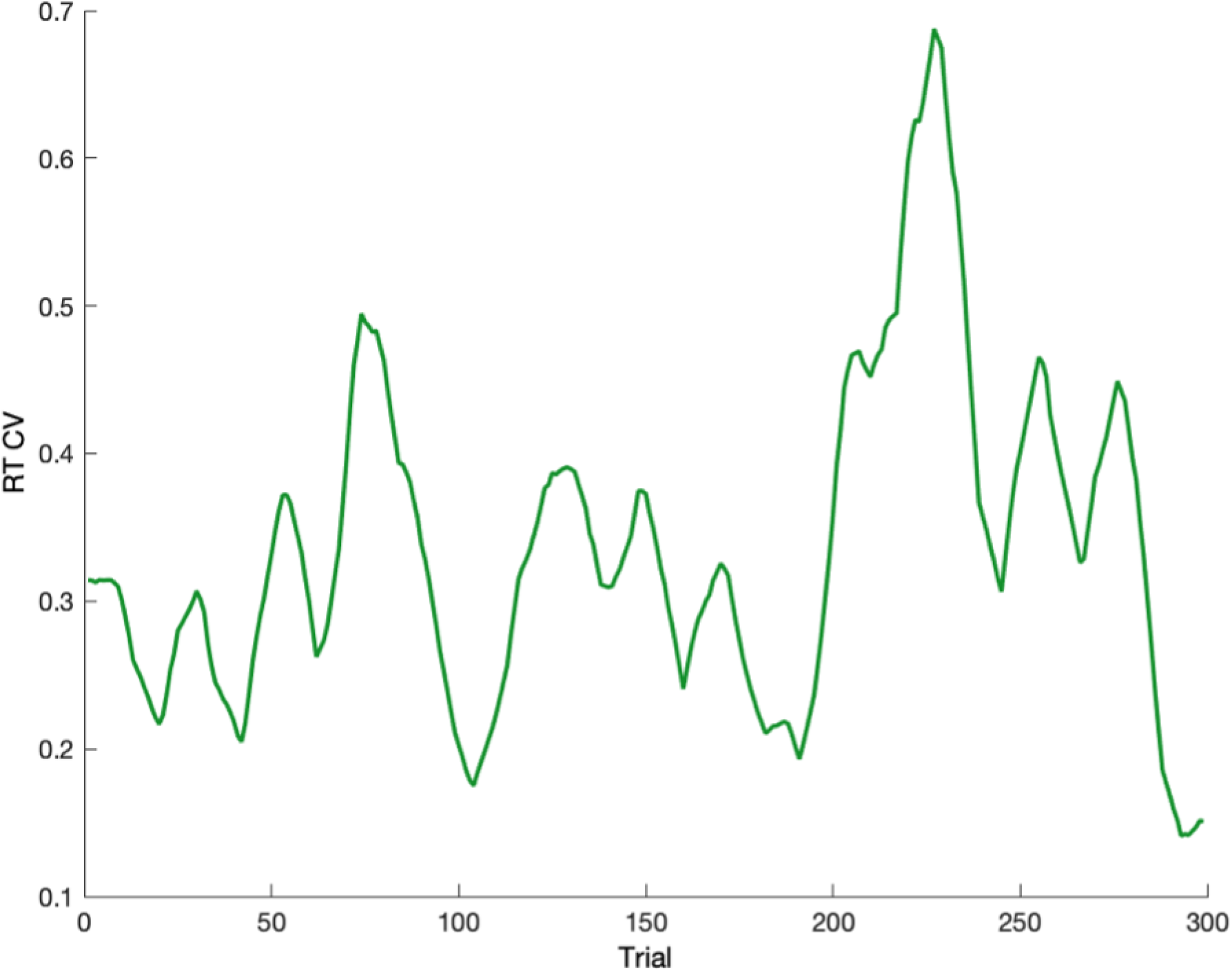
Example reaction time coefficient of variation (RT CV) of one representative participant over the course of the experiment (300 trials), indicating the variance of the 10 (log-transformed) RTs around each trial. High RT CV values indicate periods of low sustained attention.

RTs on control trials were systematically faster than on violation trials (*t*(19) = −16.46, *p* < .001). Since these two trial types appeared in random order, different degrees of trial type variability alone could lead to variance in the RT CV. To rule out that any potential effect of RT variance on ERP amplitudes might simply reflect an effect of trial type variability (i.e., more mixed vs homogenous trial types within a given time period), we regressed out trial type variability from the RT CV line vector. We did so by coding violation and control conditions as −1 and +1, respectively, and then calculating a moving average of these conditions using the same window size as the moving RT coefficient of variation (10 trials, as well as exploratory windows of 5 and 15 trials). The resulting absolute value thus indicated the degree of trial type variability in the window around each trial (bound between 0, indexing heterogenous trial types and 1, indexing homogenous trial types). We linearly regressed the RT CV against this trial type variability and took the residual variance as the final “RT CV” line.

As a result, each trial was associated with a single RT variance value, indicating the degree of homogeneity of RTs in the surrounding trials. To get a more reliable measure of sustained attention over the entire task, the RT CV was calculated over both control and violation trials. However, for subsequent ERP analyses, we were primarily interested in violation trials (basic comparisons between ERPs for violation versus control trials are reported in SBS). Therefore, we kept only RT variance values for violation trials.

### EEG Data Preprocessing

EEG data were preprocessed in Matlab R2020a using the EEGLAB (Delorme & Makeig, 2004) toolbox. Data were downsampled to 100 Hz and re-referenced to the average of the left and right mastoids. Ocular artifacts were removed using independent component analysis (Infomax ICA; Jung et al., 2001; Makeig et al., 1997) on segments spanning −500 ms pre- and 2000 ms post stimulus onset, which were extracted from continuous data filtered with a 1-30 Hz Butterworth filter. Independent components (IC) were removed which the IClabel plug-in (Pion-Tonachini et al., 2019) identified to be eye related with a probability greater than 30%. Upon visual inspection, we additionally identified and removed one channel noise IC in seven participants (see also SBS).

The corrected, continuous data were high-pass filtered (0.1 Hz, two-pass Butterworth with a 12 dB/oct roll-off) and low-pass filtered (30 Hz, two-pass Butterworth with a 24 dB/oct roll-off). Data were then epoched from −200 to 1100 ms time-locked to the critical phrase onset and baseline-corrected relative to a 200ms interval preceding the onset. Epochs with absolute values exceeding 75 *µ*V within the seven channels of interest (see below) were removed.

For analyses, we extracted epochs from trials with either semantic or morphosyntactic violations. Conservatively, only data from trials in which participants responded correctly and with RT values > 0 remained for analyses. On average, each participant contributed 129 trials (range = 114-140), of which were 62 morphosyntactic and 67 semantic violations. Data for both components were analyzed for a parietal region of interest (ROI: CP1, CP2, P3, Pz, P4, PO3, PO4). The ROI was selected to capture both the somewhat more central distribution of the N400 (e.g., Hodapp & Rabovsky, 2021) and the somewhat more posterior distribution of the P600 (e.g., Kuperberg et al., 2020; Münte et al., 1998; Tanner et al., 2017) and is an area where both effects in SBS were maximal (p. A14; see also Figure A1 in the Appendix). The components were analyzed within a 300-500 ms (N400) and 600-900 ms (P600) time window. Note that although we calculated the RT variance for each trial using a trial window around that trial, we did not average ERP amplitudes over the respective trial window. Instead, each single trial ERP amplitude was associated with a particular RT variance value calculated as described above.

### Statistical analyses

We performed linear mixed-effects models (LMM) using the package lme4 as implemented in R (R core Team, 2018) for our two planned analyses, which investigated the effect of RT variance separately on the amplitude of the N400 and P600 component on a single-trial basis. For both analyses, we controlled for (log-transformed) trial RT and violation type (semantic vs morphosyntactic) by adding them as additional fixed effects to the models. Following the recommendations by Barr et al. (2013), we tried to fit the maximal random effect structure as justified by the design but reduced its complexity successively until the model converged. Both models converged with random intercept and slope for RT variance by participant, but the N400 lacked the correlation between the random effects. The final models were thus: *P600 amplitude ~ RT variance + RT + violation type + (1 + RT variance* | *participant)* and *N400 amplitude ~ RT variance + RT + violation type + (1 + RT variance* || *participant)*. Sum coding (Schad et al., 2018) was used as contrasts for the categorical predictor violation type (morphosyntactic violation: −0.5, semantic violation: 0.5). The significance of fixed effects was determined via likelihood ratio tests to compare the fit of the model to that of a reduced model lacking the respective fixed effect but including all remaining fixed effects as well as the same random effect structure. Note that we initially modelled RT variance as a dichotomous predictor using the median split across all RTCV values within a participant (method analogous to e.g., Estermann et al., 2013). This analysis pipeline yielded identical results and we report these model outcomes in the Appendix for full transparency (Table A1). However, to avoid information loss through dichotomization (Cohen, 1983), we now model RT variance as a continuous predictor instead.

## Results

ERP means on morphosyntactic and semantic violations for high vs. low RT variance trials are shown in Figure 2. This dichotomization into high (above median) and low (below median) RT variance is for visualization only, as RT variance was modelled as a continuous predictor. As to be expected from previous literature and the results by SBS, both semantic and morphosyntactic violations generally elicited a large positive deflection approximately 600-1000 ms after phrase onset, in comparison to correct control phrases. In contrast, only semantic violations exhibited an increase in N400 amplitudes. Still, in order to increase power and be able to compare results between components, we included data from both violation types in the models testing the effect of RT variance on both P600 and N400 amplitudes.

**Figure 2.**
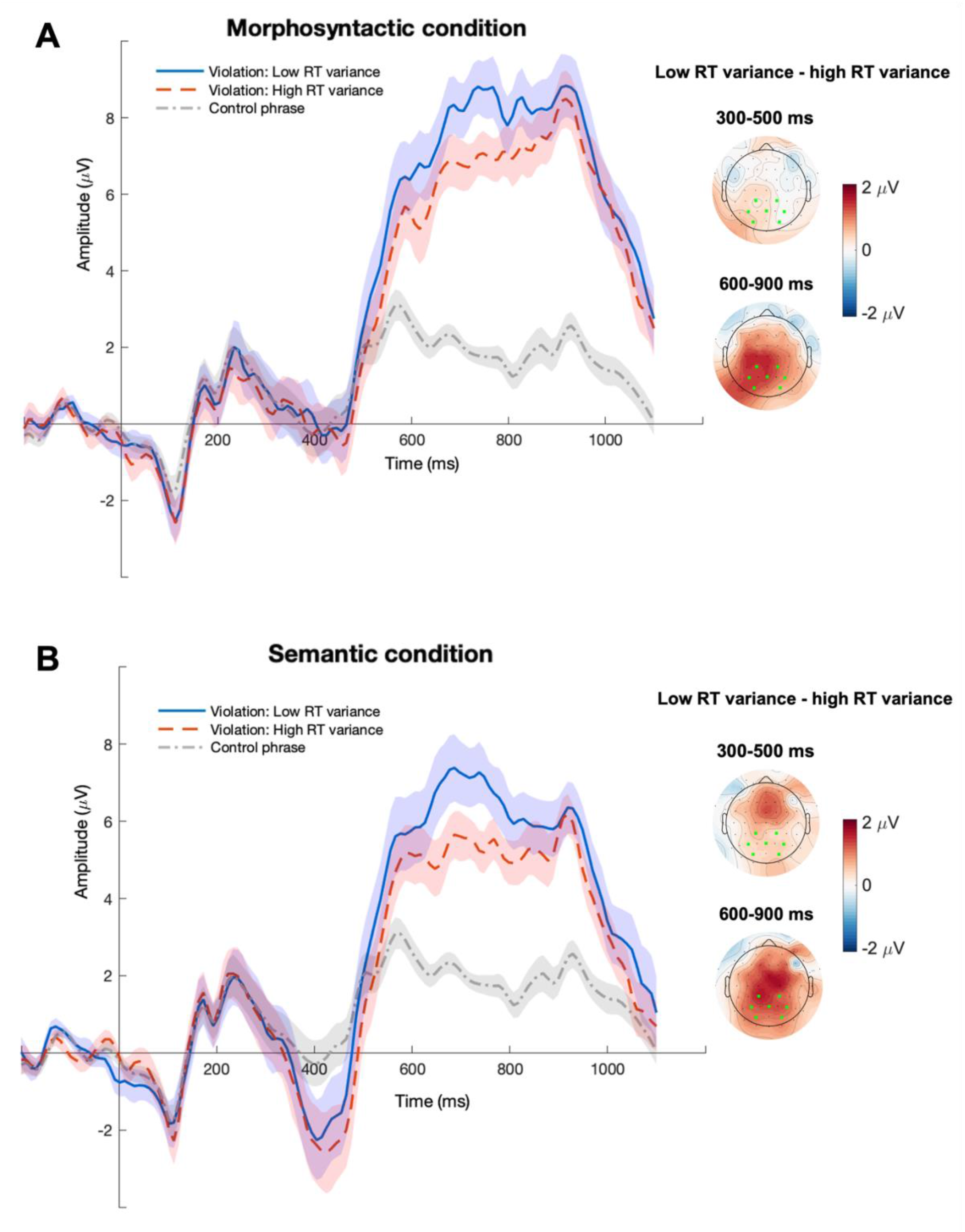
Grand average waveforms of trials with morphosyntactic violations (A) and semantic violations (B) at parietal ROI in high vs. low RT variance periods, time-locked to the onset of the violation phrase. The additional control phrase mean includes data from all non-violation targets (on average 692 phrases per participant, range = 663-715). Control phrase means are for visualization only, that is, they were not part of our statistical analysis. Error bands indicate the SEM. The topography of the RT variance effect (high vs. low) within the N400 and P600 time window is plotted to the right of the waveforms. Note that trial outliers (values exceeding +/-75 µV) in channels outside the ROI (green squares) were additionally excluded here for illustration only. Note also that the categorization into high (above median) vs low (below median) used in these plots is for visualization only, as RT variance was modelled as a continuous predictor in our models.

### P600

P600 amplitudes were significantly larger in trials with morphosyntactic than semantic violations (*β* = 2.01, *SE* = 0.33, *t* =6.18, *χ^2^* = 37.86, *p* < .001; Fig 3A, left), in line with previous evidence that P600 amplitudes are primarily (though not exclusively) sensitive to syntactic violations (e.g., Osterhout & Mobley, 1995). Further, there was a negative effect of RT on the P600, thus amplitudes were smaller in trials with longer RTs (*β* = −4.72, *SE* = 0.46, *t* = −10.37, *χ^2^* = 104.44, *p* < .001; Fig 3B, left). Crucially, the model additionally revealed a significant, negative effect of RT variance on the amplitude of the P600. Hence, amplitudes were smaller in trials that had occurred during periods with higher RT variance (indexing lower sustained attention) than trials that had occurred during periods of lower RT variance (indexing higher sustained attention) (*β* = −3.18, *SE* = 1.4, *t* = −2.28, *χ^2^* = 4.58, *p* = .03; Fig 3C, left). Finally, two exploratory models demonstrate that the effect is not restricted to our chosen time window for the RT CV, as the effect of RT is also significant when it is calculated over 5 (*β* = −3.26, *SE* = 1.18, *t* = −2.77, *χ^2^* = 5.88, *p* = .02) and 15 trials (*β* = −3.78, *SE* = 1.31, *t* = −2.9, *χ^2^* = 6.71, *p* = .009)^1^.

**Figure 3.**
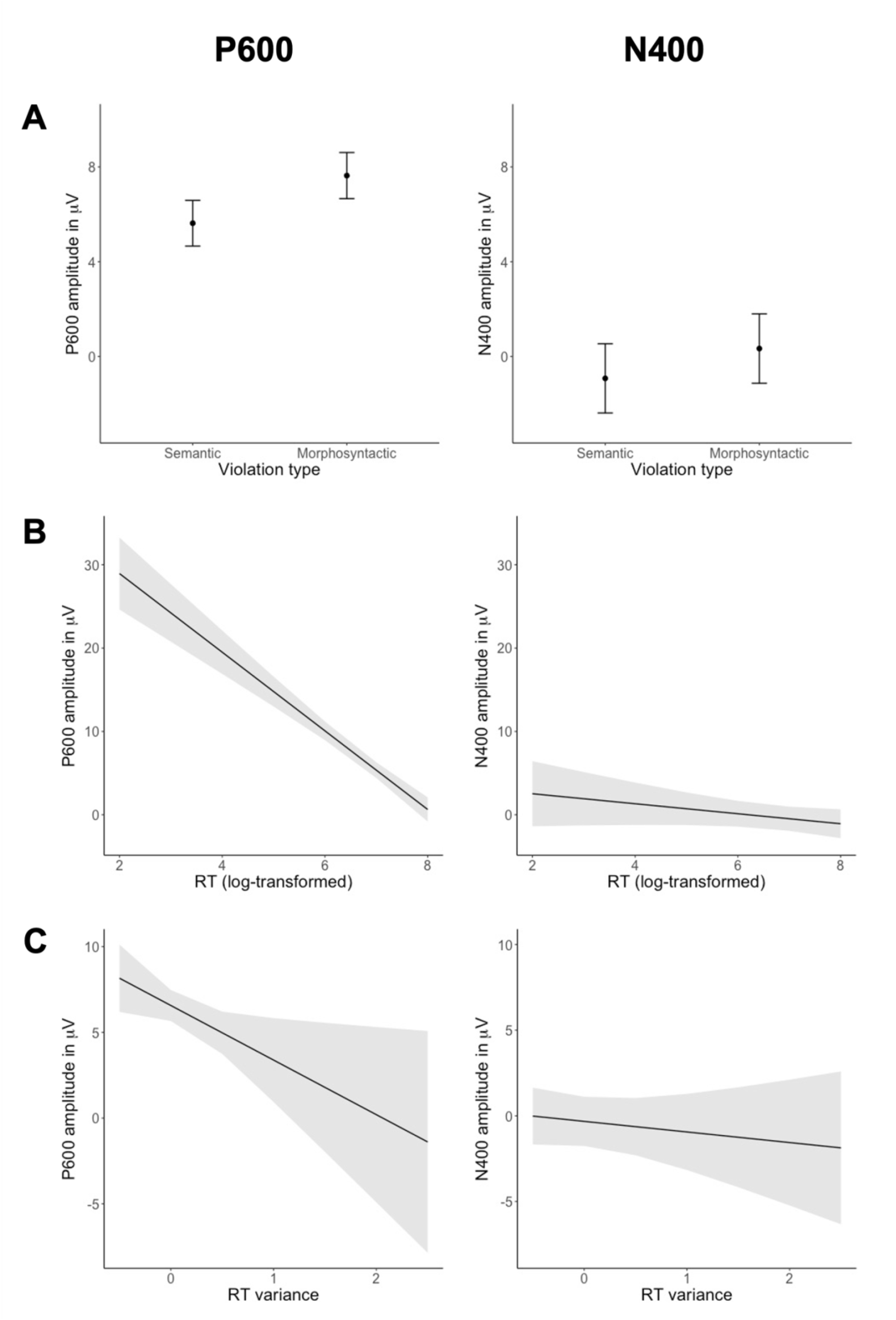
Model predictions for each of the three predictors in our model for the P600 (left) and N400 (right). Error bars indicate 95% confidence intervals. Most importantly, RT variance (C) had a negative effect on the amplitude of the P600 (left), but not N400 (right).

Note that the two RT predictors - RT of the respective trial and RT variance - might be related and thus, might also share variance in explaining the P600 amplitude. However, the variance inflation factors of our planned model did not indicate any collinearity problem (Violation type: 1.02, RT: 1.02, RT variance: 1).

### N400

N400 amplitudes were more negative on semantic than syntactic violations (*β* = 1.26, *SE* = 0.28, *t* = 4.53, *χ^2^* = 20.39, *p* < .001; Fig 3A, right) consistent with previous evidence that N400 amplitudes are sensitive to semantic but not syntactic violations (e.g., Kutas & Federmeier, 2011). However, N400 amplitudes were neither affected by RT (*β* = −0.60, *SE* = 0.39, *t* = −1.53, *χ^2^* = 2.3, *p* = .13; Fig 3B, right), nor RT variance (*β* = −0.62, *SE* = 0.86, *t* = −0.72, *χ^2^* = 0.57, *p* = .45; Fig 3C, right).

In order to make sure we did not miss a potential effect of RT variance on the N400, we exploratorily conducted three additional analyses. For better comparison between the results regarding the two components, the planned N400 model included data from both semantic and morphosyntactic violations. However, the N400 is primarily modulated by semantic factors and indeed, Figure 2A also indicates that there was no N400 effect for morphosyntactic violations to begin with. To take this into account, we fit an additional N400 model including only data from semantic violations (and thus also excluding the fixed factor of violation type). However, even with semantic violations only, N400 amplitudes were not modulated by RT variance (*β* = 0.42, *SE* = 1.29, *t* = 0.33, *χ^2^* = 0.11, *p* = .75). Likewise, for better comparison, we used the same spatial ROI for both components (parietal cluster around Pz), although the N400 is often observed at slightly more central area (e.g., Hodapp & Rabovsky, 2021; see also Fig. A2 in the Appendix). Thus, we also checked whether we missed a potential effect of RT variance in the 300-500 ms time segment at a centro-parietal channel cluster (C1, C2, CP3, CPz, CP4, P1, P4). However, the effect of RT variance remained non-significant and the direction of the estimate was – just like the one in the planned model above – even in the opposite direction (*β* = −0.4, *SE* = 0.95, *t* = −0.42, *χ^2^* = 0.23, *p* = .63). Finally, we again made sure that the results are not contingent on the specific time window for calculating the RT CV. Both models with alternative time windows also failed to find an effect of RT variance on the N400 (5 trials: *β* = −0.29, *SE* = 0.6, *t* = −0.48, *χ^2^* = 0.29, *p* = .59; 15 trials: *β* = −0.57, *SE* = 1.04, *t* = −0.55, *χ^2^* = 0.35, *p* = .55).^2^

### Comparison of RT variance effect on P600 and N400

Lastly, we directly compared the amount of RT variance explained in each of the ERP amplitudes. To do so, we built a regression model of ERP amplitude which included not only the predictors violation type, single trial RT, and RT variance, but also the interaction between RT variance and ERP component (P600 vs N400). Note that the P600 and N400 have opposing polarity: More positive amplitudes signify a larger P600 but smaller N400, and vice versa. In order to directly compare the size of the effect of RT variance on each component in this joined model, we therefore reversed the sign of the amplitude values in the N400 time window, so that larger values also reflect a larger (more negative) N400. ERP component was sum coded (N400 = −.5, P600 = .5) and due to its increased complexity, the model converged only with random intercepts by subject. In this comparison model, the interaction between RT variance and component was significant (*β* = −2.86, *SE* = 1.32, *t* = −2.17, *χ^2^* = 4.72, *p* = .03): RT variance negatively affected P600 amplitudes (*β* = −2.37, *SE* = 0.93) but not N400 amplitudes (*β* = 0.48, *SE* = 0.93).

## Discussion

To investigate whether the P600 and N400 might be dissociated on the automaticity vs control dimension, we tested if they are differentially modulated by sustained attention as indexed by RT variability. Indeed, P600 amplitudes were significantly affected by RT variance in that they were larger with lower RT variability (i.e., the higher sustained attention). This effect emerged even when controlling for individual trial RT and considering only trials in which participants responded correctly. In contrast, the amplitude of the N400 was not affected by RT variability. These results suggest that the P600 component is sensitive to the current level of sustained attention while the N400 component is less so. Since sustained attention is needed to enable executive resources needed for controlled processes, these findings thus provide further evidence that P600 amplitudes reflect more controlled and N400 amplitudes more automatic aspects of sentence comprehension (Rabovsky & McClelland, 2020).

Our findings also align well with previous evidence on differential effects of task relevance on the two components (e.g., Schacht et al., 2014), together fostering the idea that attention is a strong modulator of the P600, but not the N400. In contrast to previous studies, the current study did not manipulate the relevance of the stimuli, or the instruction given to participants, but rather measured sustained attention more directly, across the experiment, using reaction time variability as an index. This measure also takes into account that sustained attention naturally fluctuates over time. To our knowledge, this was the first attempt to adopt this method from fMRI research on inhibition and attention (e.g., Esterman et al., 2013; Fortenbaugh et al., 2018) in the context of (language-related) ERPs. Evidently, the task demands in the commonly used continuous performance paradigms – execution or inhibition of quick consecutive responses to simple perceptual differences – differ from a linguistic judgement task as used here. Our results indicate that RT variability might have further potential as a measure for continuous monitoring of fluctuations in sustained attention in language processing, complementing recent related indices such as the shape of RT distributions in self-paced reading (e.g., Payne & Federmeier, 2017).

In the current study, participant actively judged whether sentences were anomalous or not, providing reaction times which could be used to calculate RT variance as an index for sustained attention. The P600 has also been observed in paradigms without an overt task (Hagoort et al., 1993; Kaan et al., 2000). Evidently, our approach of measuring sustained attention in these paradigms would not be possible and the present finding might not inform us about the controlled vs automatic nature of the P600 in these passive paradigms. Thus, strictly speaking, the current findings cannot disambiguate whether sustained attention indeed affects the P600 which also appears in passive linguistic tasks, or rather a separate, more domain-general late positivity that is decision-related and tied to a response. However, a lot of what we know so far about the P600 originates from many studies where participants were also required to make judgements about the acceptability or interpretation of the respective sentences (Allen et al., 2003; Gunter & Friederici, 1999; Hahne & Friederici, 1999; Kaan & Swaab, 2003; Kim & Osterhout, 2005; Osterhout & Mobley, 1995, just to name a few). The current findings are thus relevant for a large portion of the literature on the P600 and the effect of RT variance on the P600 should potentially also be observed in such paradigms, even when the judgement is prompted after the end of the sentence, and not only – as in the present experiment - as soon as the anomaly is detected.

We additionally found that P600 - but not N400 - amplitudes exhibit a negative relationship to RT, which is also in line with the assumption that the P600 is more tightly linked to performance and executive control than the N400. Note though, that we quantified P600 amplitude as the mean amplitude in a pre-defined time window and SBS’s study suggests that the P600 latency is RT-rather than stimulus onset-aligned. Thus, our negative relationship might be an inevitable consequence: The longer the RT, the more the P600 “moves away” from the pre-defined window, decreasing the mean amplitude in this window. However, SBS also found such a negative correlation between P600 and RT when quantifying the P600 amplitude as the 60ms around the component’s peak (see their supplementary material). Though pending more direct investigation, the observed relationship between RT and ERP amplitude might indicate that the amplitude of the positivity on a given sentence reflects how much executive resources (such as reorientation of attention) are recruited upon detecting the violation and in turn, how fast the reader can respond to it.

As predicted, the amplitude of the N400 was not significantly affected by RT variability, indicating this earlier component is less reliant on sustained attention and might thus need less or no cognitive control. It is possible that also the N400 is modulated by sustained attention, but to a much smaller degree. However, the non-significant effect observed on the N400 even goes into the opposite direction, with less negative amplitudes in periods of *high* sustained attention. Additionally, we demonstrated that the null effect on the N400 is neither contingent on the type of violation, nor the channel cluster, nor the time window considered for RT variance. Together with literature suggesting that the N400 is even present when the participant’s attention is drawn away from sentence processing (Schacht et al., 2014) and during unconscious priming (Kiefer, 2002, van Gaal et al., 2014), the present results support the notion that the N400 indexes semantic processes that occur rather automatically.

Our finding that sustained attention modulated P600 amplitudes also has interesting implications for a current hypothesis on the neural generator of the P600. Sustained attention is assumed to be enabled by arousal, possibly via tonic norepinephrine (NE) levels (Esterman & Rothlein, 2019; Van Den Brink et al., 2016). Under medium tonic NE levels and an optimal state of arousal, sustained attention on the current task is high. When tonic NE levels and arousal are very low or too high, participants are either drowsy or highly alert and sustained attention on the current task is compromised. Interestingly, phasic noradrenaline activity also exhibits such a non-linear relationship with tonic NE levels: Strong responses under medium tonic levels, small or absent responses under very low or high tonic levels (Aston-Jones & Cohen, 2005). Crucially, the P600, just like the P3, has been proposed to reflect such phasic NE release (Nieuwenhuis et al., 2005; Sassenhagen et al., 2014; Sassenhagen & Bornkessel-Schlesewsky, 2015). Taken together, our data might contribute to the assumption that under optimal arousal (medium NE tonic levels), sustained attention on the task is high, and participants have executive resources available to detect and/or “act upon” violations, which leads to good performance and a large P600. Conversely, under very low or very high arousal and tonic NE levels, sustained attention suffers, so participants are disengaged, are less likely to detect violations, diminishing the P600 and leading to poor performance. Indeed, in periods of high RT variance, accuracy in violation trials was significantly lower than in periods of low RT variance (*β* = −2.69, *SE* = 0.59, *t* = −4.58, *χ^2^* = 15.39, *p* < .001)^3^. Although this post-hoc accuracy analysis was not our main focus here, this might imply that low sustained attention reduces the ability to detect linguistic violations in the first place. Importantly however, we found the effect of RT variance on the P600 even though we considered only correct violation trials in our analysis. This might suggest that sustained attention affects processing aspects underlying the component over and above error detection, possibly involving controlled revision and behavioral adaptation, akin to the proposed “network reset” induced by phasic norepinephrine (Bouret & Sara, 2005).

In conclusion, the current study suggests that the P600 is sensitive to sustained attention, providing further evidence that the underlying process might require cognitive control. In turn, the N400 seems to depend less on sustained attention, further indicating that it reflects more automatic aspects during sentence comprehension. Our adopted measure of reaction time variability could be a promising method to monitor the impact of ongoing fluctuations in sustained attention on higher level cognitive processing such as language comprehension.

## Data availability statement

All analyses scripts and final data are accessible on OSF (https://osf.io/xedyq/).

## Author contribution

Conceptualization: FC, MR, IW; Methodology, Visualization, formal Analysis: FC supervised by MR; Writing - Original Draft Preparation: FC supervised by MR; Writing - Review & Editing: FC, MW, IW, MR; Supervision: MR, MW, IW; Funding Acquisition: MR, MW, IW.

## Appendix

**Table A1.**
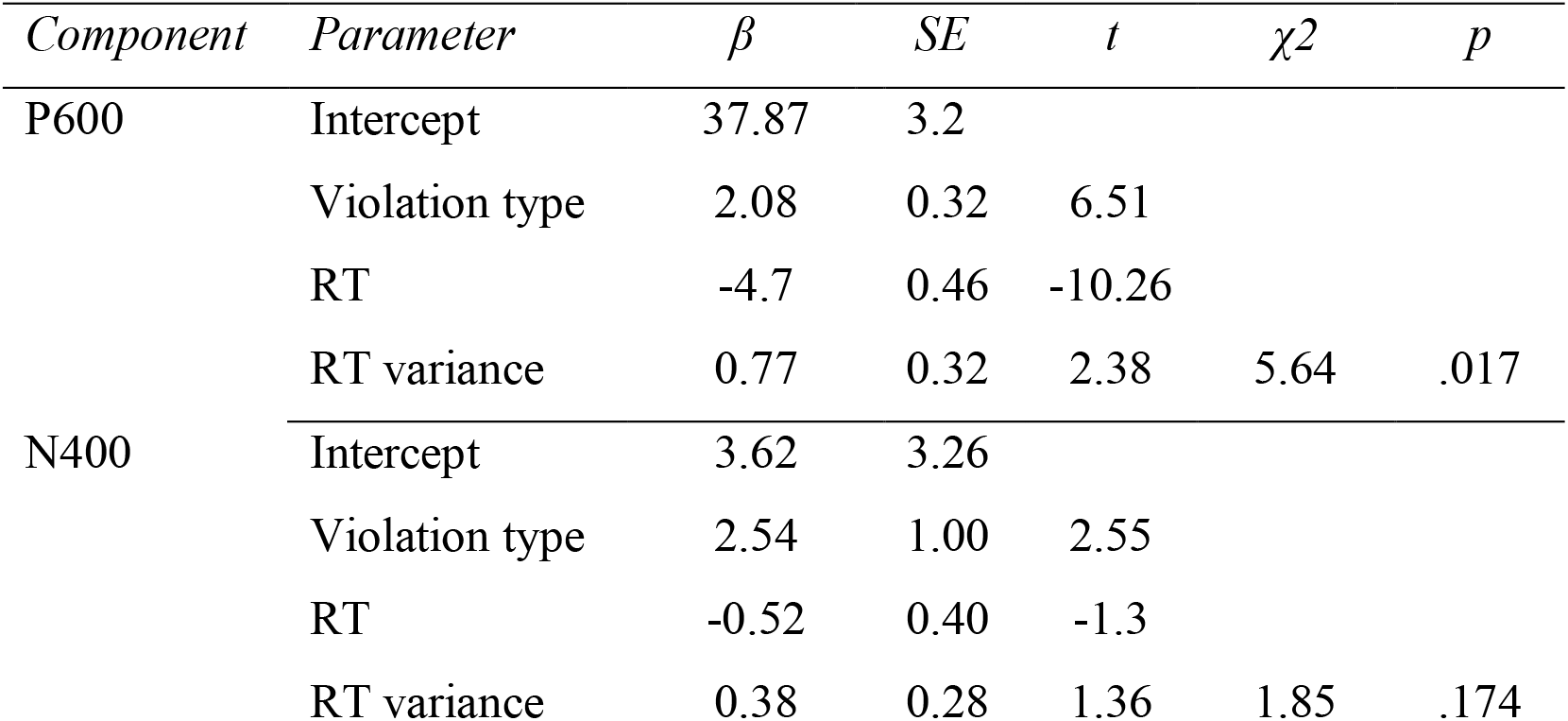
Model outputs from our original analyses in which RT variance was modeled as a binary predictor split into low RT variance (high sustained attention) vs high RT variance (low sustained attention) based on median split across all RT CV values within each participant. RT variance is sum coded (high RT variance = −.5, low RT variance =.5). χ^2^ and p values for RT variance are derived from a likelihood ratio test against a model lacking the respective predictor.

**Table A2.**
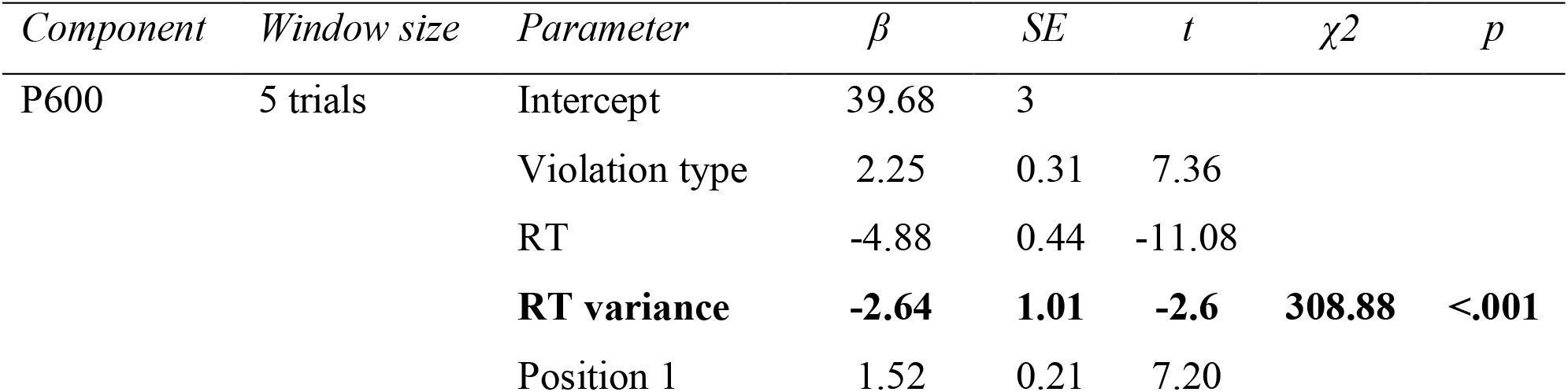
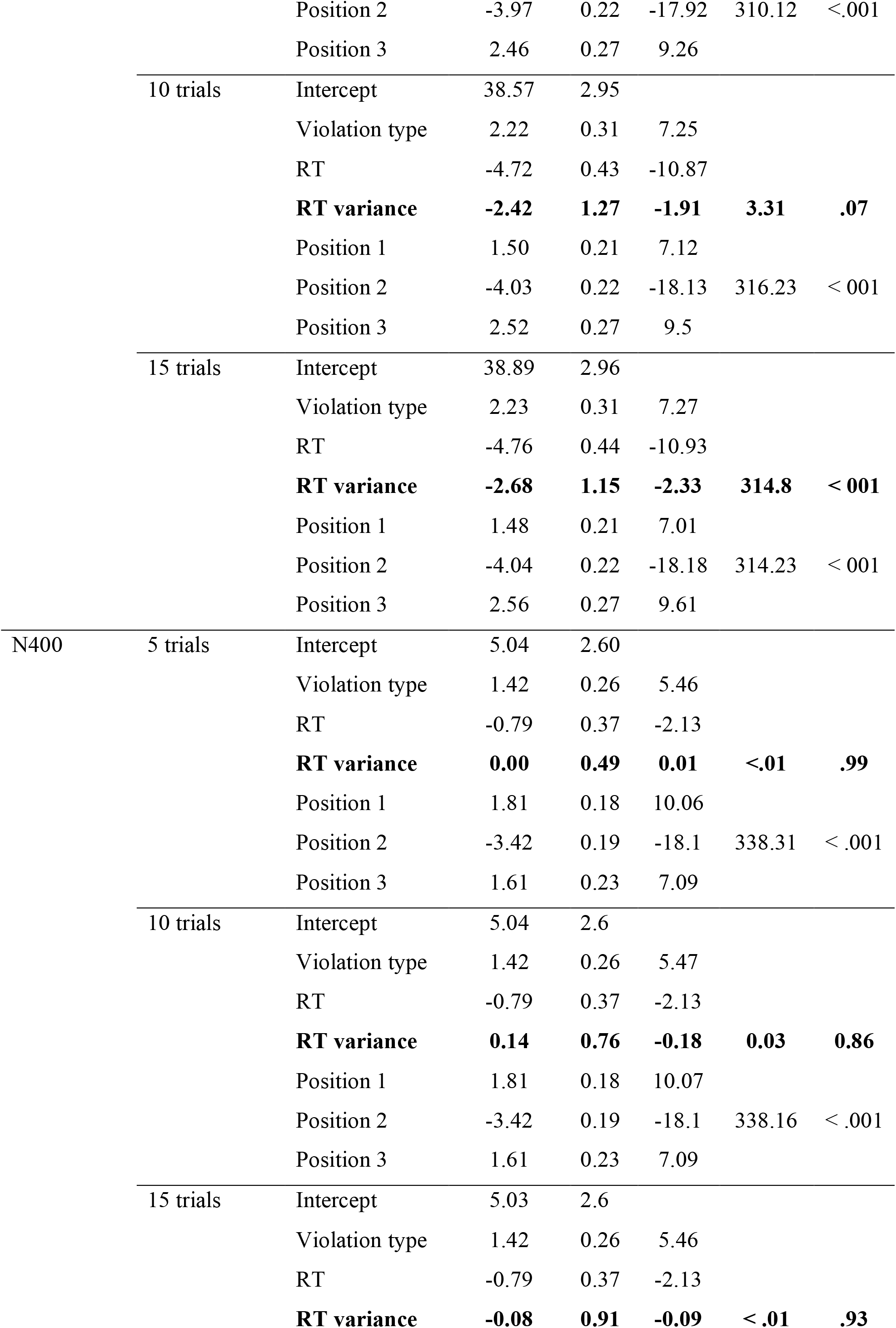
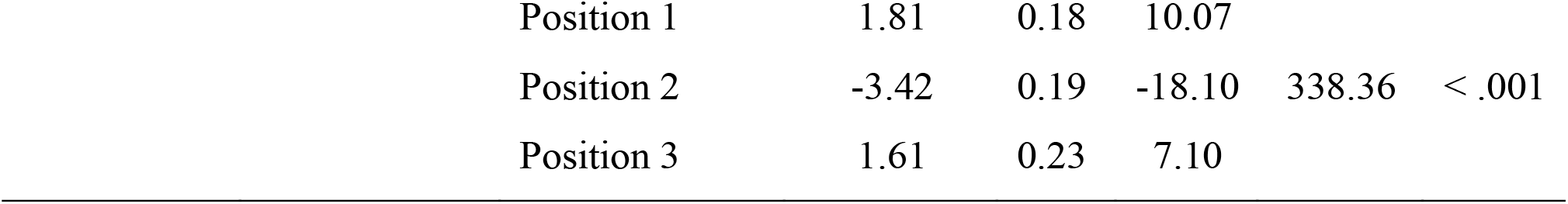
Model outputs for the P600 and N400 models including RT variance across the three time windows (5, 10, and 15 trials) and additionally including position as a nuisance predictor (sum coded). χ^2^ and p values for the predictors of interest (RT variance, position) are derived from a likelihood ratio test against a model lacking the respective predictor.

**Figure A1.**
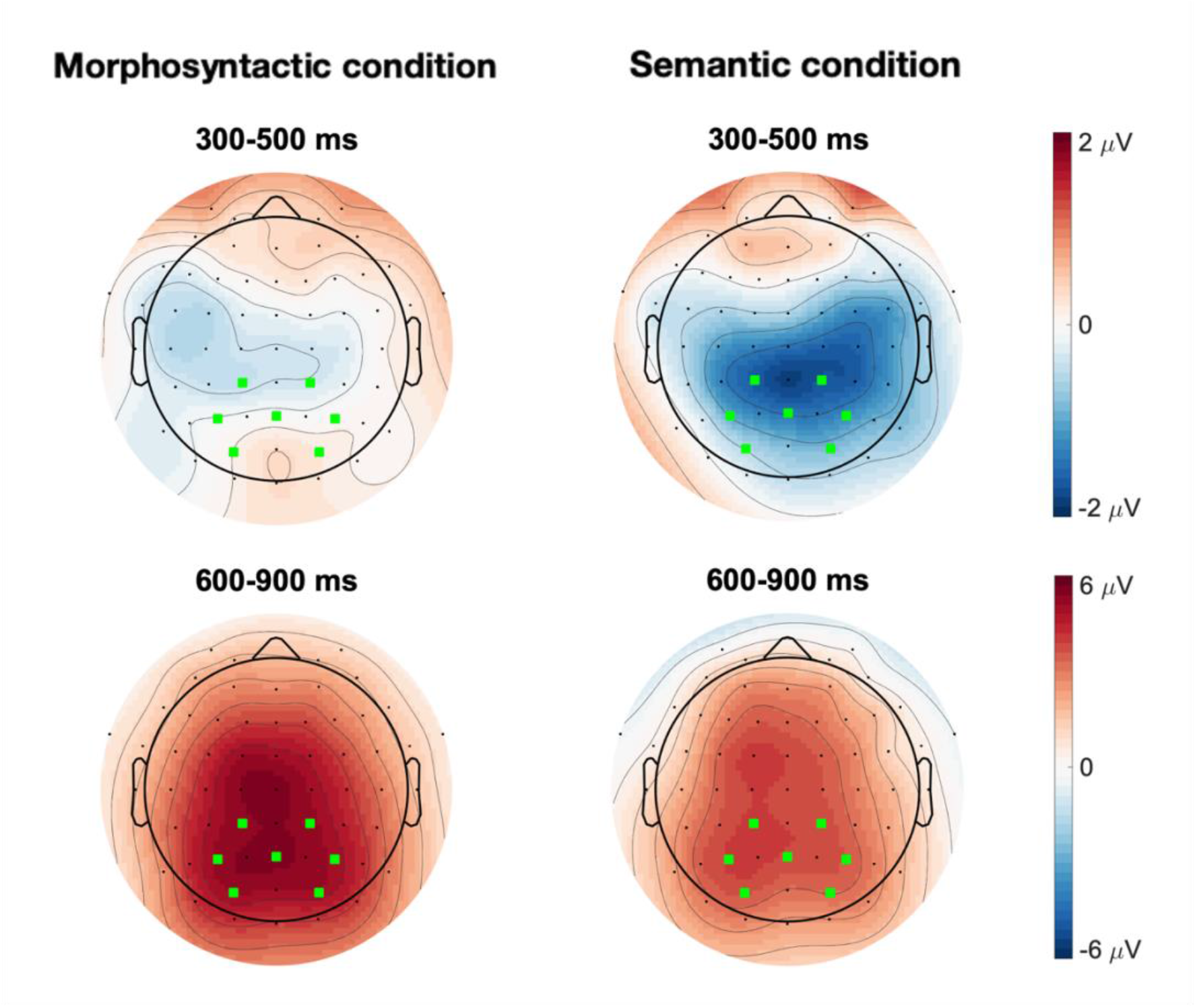
Topography the effect of morphosyntactic violations minus control (left) and semantic violations minus control (right). On the top is the early time window (N400) and bottom the later time window for the (P600). Green squares mark our centro-parietal ROI for the N400 and P600.

## Footnotes

1 Additionally, we explored the influence of position of the respective noun, by including it in each of the three models. As indicated in Table A2 (Appendix), the P600 was generally smaller on the second noun than on the first or third noun. RT variance remained a significant predictor of P600 amplitudes, two-sided in both time windows 5 and 15 (both *p* < 001) and in a one-sided test with time window 10 (*p* = .07). Including position possibly weakened the effect of RT variance since position showed the opposite effect on RT variance than position on the P600 (i.e., larger RT variance on the second noun, *β* = 0.01, *SE* = 0.00, *t* = 2.3, *p* = .02; and smaller RT variance on the third noun, *β* = −0.01, *SE* = 0.01, *t* = −2.44, *p* = .01; each compared to the grand mean). That the effect of position was the same for the N400 (Table A2) indicates the EEG amplitude was generally smaller in position two and was thus, generally negatively correlated with RT variance. Future research using paradigms without different positions would be desirable in order to avoid this confound.

2 AAgain, we also explored the influence of position of the respective noun, by including it in the models (for the output, see table A2 in the Appendix). The N400 was also more negative on the second noun than on the first or third noun, indicating large portions of the ERP amplitude in general showed this effect (i.e., not specific to either of the components). All models failed to detect an effect of RT variance on the N400.

3 A Retrieved from a mixed logistic regression model in which accuracy was predicted by RT variance and violation type (as a nuisance predictor), incl. random intercept and slope for RT variance by participant.

